# Chromatin interactions and expression quantitative trait loci reveal genetic drivers of multimorbidities

**DOI:** 10.1101/340216

**Authors:** Tayaza Fadason, William Schierding, Thomas Lumley, Justin M. O’Sullivan

## Abstract

Clinical studies of non-communicable diseases identify multimorbidities that reflect our relatively limited fixed metabolic capacity. Despite the fact that we have ∼24000 genes, we do not understand the genetic pathways that contribute to the development of multimorbid non-communicable disease. We created a “multimorbidity atlas” of traits based on pleiotropy of spatially regulated genes using convex biclustering. Using chromatin interaction and expression Quantitative Trait Loci (eQTL) data, we analysed 20,782 variants (p < 5 × 10^−6^) associated with 1,351 phenotypes, to identify 16,248 putative eQTL-eGene pairs that are involved in 76,013 short- and long-range regulatory interactions (*FDR* < 0.05) in different human tissues. Convex biclustering of eGenes that are shared between phenotypes identified complex inter-relationships between nominally different phenotype associated SNPs. Notably, the loci at the centre of these inter-relationships were subject to complex tissue and disease specific regulatory effects. The largest cluster, 40 phenotypes that are related to fat and lipid metabolism, inflammatory disorders, and cancers, is centred on the *FADS1-FADS3* locus (chromosome 11). Our novel approach enables the simultaneous elucidation of variant interactions with genes that are drivers of multimorbidity and those that contribute to unique phenotype associated characteristics.

## Introduction

The rising incidences of complex diseases and there accompanying cost burdens^1,2^ are driving a shift in disease research and management from the single-disease to a broader paradigm that accommodates a patient’s overall health^2,3^. Completing this paradigm shift requires many advances including a greater understanding of the genetic aetiology of the observed multimorbidities, which remain largely unknown^4^. The importance of understanding the genetic aetiology of multimorbidity is further compounded by the fact that almost 90% of single-nucleotide variations (SNPs)—a major source of shared heritability of polygenic disorders—do not fall within coding regions of the genome^5^.

Cross-phenotype genetic studies have been conducted on a small number of complex traits (the largest one studied 42 traits^6^) with known associations using methods that included systematic reviews^7^, LD score regression^8,9^, polygenic risk scores^10^, Probabilistic Identification of Causal SNPs^11^ and Bayesian colocalisation tests^6^ on Genome Wide Association Study (GWAS) summary or molecular data. These studies^6–8,10,11^ typically use SNPs, or the genes that are nearest to—or in LD with—the SNPs as the putative genetic drivers of cross-phenotype associations. These approaches are limiting as evidence increasingly shows that gene regulatory elements (*e.g*. enhancers) can impact distant genes more strongly than the genes harbouring, or closest to, them as a result of physical interactions with distal chromatin regions within the 3D organization of the genome^12–18^.

In this study, we undertook a discovery-based approach to identify genes that are spatially associated with eQTLs from phenotype-associated SNPs for all human traits within the GWAS Catalog. We then used a convex biclustering algorithm to identify multimorbidities by clustering the traits according to shared target genes. Lastly, we demonstrate the utility of this approach in understanding the common genetic aetiology of multimorbid traits.

## Results

### GWAS SNPs mark spatial regulatory regions

Disease-associated SNPs often mark gene enhancers, silencers and insulators^19,20^. We set out to identify the genes whose transcript levels are associated with regulatory regions marked by disease and phenotype associated SNPs (daSNPs). We downloaded 20,782 unique daSNPs (p<5 × 10^−7^) from the GWAS Catalog (www.ebi.ac.uk/gwas/). The CoDeS3D pipeline^14^ was used to interrogate Hi-C chromatin interaction libraries^21^ to identify genes that are captured as physically interacting with the GWAS SNP-labelled regions within nuclei from one or more of seven cell lines (GM12878, HMEC, HUVEC, IMR90, K562, KBM7 and NHEK) ^21^. The resulting 1,183,037 spatial SNP-gene pairs were used to query the GTEx database (www.qtexportal.orq, multi-tissue eQTLs analysis v4) to identify eQTL-eGene pairs.

A total of 7,776 (∼38.4%) of the GWAS SNPs analysed were associated with a change in the expression (*i.e*. eQTLs) of 7,917 eGenes at an *FDR* ≤ 0.05 (Benjamini Hotchberg^14^), for a total of 16,248 distinct eQTL-eGene pairs (76,013 interactions in different tissues). daSNPs with significant eQTLs are distributed (range = 86 – 764) across chromosomes 1 -21. The distributions of daSNPs and eQTLs correlate (Pearson’s *r*, 0.86 and 0.74 respectively) with the sizes of the chromosomes, with Chromosomes 1 and 21 having the most and least eQTLs respectively (a in S1 Fig). Similarly, the number of eGenes on chromosomes correlated (Pearson’s *r* = 0.86) with the number of genes per chromosome (b in S1 Fig). Notably, none of the 182 daSNPs on chromosome X were identified as having significant spatial eQTL effects with any gene (b in S1 Fig). However, 41 genes on the X chromosome are affected by *trans*-eQTLs from other chromosomes (b and d in S1 Fig). The underrepresentation of GWAS SNPs and eQTLs on the X chromosome can be explained by the exclusion of X-linked genetic variants from GTEx and two-thirds of GWAS^22,23^. As none of the databases referenced here have Y or mitochondrial data, there are also no SNPs nor eGenes identified on those chromosomes.

Only 24.3% of the eQTL-eGene pairs were correctly mapped in the GWAS Catalogue and these contribute 18.5% of the tissue specific eQTL-eGene interactions (e in S1 Fig). This finding is consistent with previous observations^14,23^ that highlighted the discordant results between the ‘nearest gene’ and eQTL-based assignment of GWAS SNPs to target genes. Of the 7,917 affected eGenes, 70.0% (5,545) were associated with only *cis*-spatial interactions (*i.e*. both partners are from the same chromosome and separated by <1,000,000 bp), 19.3% (1,528) by only *trans*-interactions (*i.e*. both partners are from the same chromosome and separated by >1,000,000 bp, or from different chromosomes), and 10.7% (844) by both *cis*and *trans*-interactions (e in S1 Fig, S2 Fig, S1 Table). The most significant eQTL-eGene interactions involved associations between: 1) the long intergenic non-protein coding RNA *CRHR1-IT1* (Chr. 17) and rs12373124, rs12185268, rs1981997, rs2942168, rs17649553, rs17689882 and rs8072451 in subcutaneous adipose tissue (*p*_eQTL_ range, 1.39 × 10^−95^ – 4.33 × 10^−94^); and 2) *PEX6* (Chr. 6) and rs9296404 in skeletal muscle (*p*_eQTL_ 1.59×10^−93^; c and d in S1 Fig).

### Multimorbid phenotypes cluster around shared eGenes

We reasoned that the common pathogenesis seen in polygenic diseases and phenotypes occurs because they share common biochemical pathways and genes. To identify associations among phenotypes, we populated phenotype matrices according to their common eQTLs or eGenes (a and b in S3 Fig, S2 Table). Q-Q plots of the ratios of shared eQTLs and eGenes identified greater coverage of intermediate values between 0 and 1 that was consistent with increased information within the phenotype matrix that was populated using the eGenes (a and b in S3 Fig). To ensure that the eGene association pattern was non-random, we generated 1000 null datasets by pooling together all phenotype associated eGenes and randomly reassigning them to phenotypes, such that each control phenotype had the same number of eGenes as its corresponding test phenotype. Phenotype matrices populated by the mean null phenotype eGenes ratios had different association and distribution patterns from those generated for the test phenotypes (c in S3 Fig, S2 Table).

Of the 1,351 phenotypes analysed, 615 are significantly associated with ≥4 eGenes. Convex biclustering^24^ revealed the intricate inter-relationships among the 618phenotypes have (Fig 1). Phenotype clusters included: a) closely related measures of a phenotype (e.g. hypertension, blood pressure and pulse pressure); b) phenotypes which are observed as co-or multimorbid (*e.g*. white matter hypersensitivities, stroke and dementia^25^; or ovarian cancer, interstitial lung disease, Alzhiemer’s disease and other cognitive disorders ^26–28^); and c) phenotypes that have controversial reports of inter-phenotype association *e.g*. autism spectrum disorder and iron biomarker levels^29,30^. Notably, the observed pattern of multimorbidity derived from shared eGenes is different from the inter-relationships between phenotypes with ≥4 SNPs (S4 Fig).

**Fig 1.**
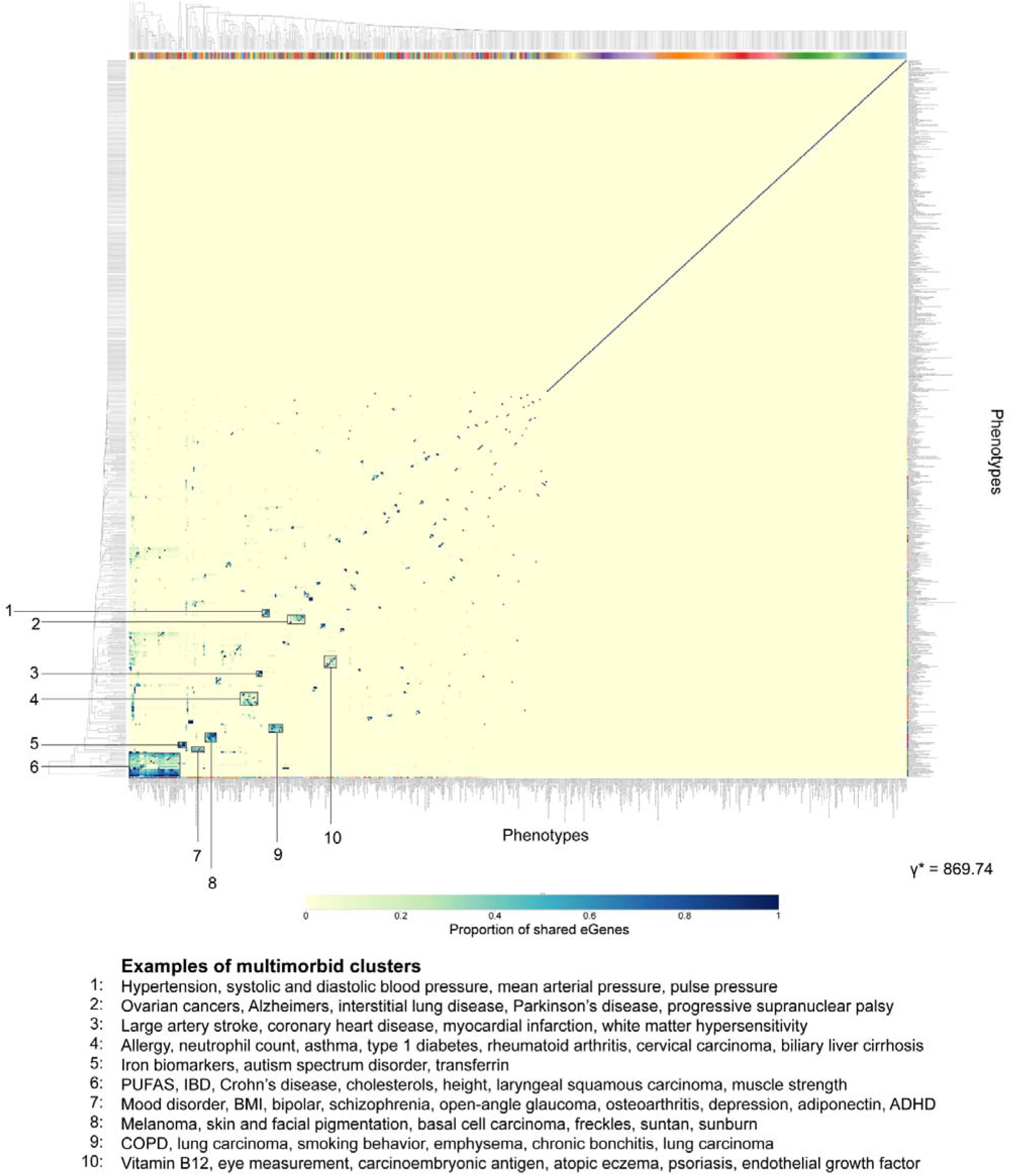
Phenotypes cluster based on common spatial eGenes. The heat map highlights the multimorbidity of 618 phenotypes that share ≥ 4 eGenes with at least one other phenotype. Unsupervised clustering was performed using convex biclustering algorithm from the *cvxbidustr* R package. Deep blue squares indicate higher proportions of shared eGenes, with 1 being the highest and indicating that two phenotypes have the same set of eGenes. Ten notable clusters are annotated and described. A complete list of inter-phenotype shared eGene proportions is presented in S2 Table.

Our discovery-based approach confirms earlier observations of pulmonary multimorbidity and genetically controlled regulatory variation in the *CHRNA* region^31^ (S6 Fig and S7 Fig). It is noteworthy that eGenes that are common to most phenotypes in the clusters lie adjacent to each other in a contiguous genomic region. This is exemplified by phenotypes that centre on: a subset of immune related disorders cluster about the *PGAP3 – GSDMA* locus (Chr 17: 37,827,375 – 38,134,431; hg19); skin pigmentation and skin cancer are clustered about the *SPATA33 – URAHP* region (Chr 16:89,724,152 – 90,114,191; hg19); while a mood disorder cluster is built about the *NT5DC2 – TMEM110* locus (Chr 3:52558385 – 52931597; hg19; S5 Fig).

The largest observed multimorbid cluster (Fig 1, cluster#6) is an outgroup of phenotypes located in the bottom left hand corner of the matrix that highlights interrelationships among polyunsaturated fatty acids (PUFAs), Crohn’s disease, inflammatory bowel disease, colorectal cancer, laryngeal squamous carcinoma, insulin sensitivity, comprehensive strength index, and cholesterol levels (Fig 2a). The cluster is built about a 283 kb region on chromosome 11 that contains the *DAGLA, MYRF, TMEM258, RP11-467L20, FADS1, FADS2, FADS3*, and *BEST1* genes (Fig 2b; S3 Table). Consistent with this observation, genetic variations in the *FADS1 – FADS3* region have previously been associated with alterations in the synthesis of PUFAs^32^, inflammatory bowel diseases ^33^, cholesterol levels and BMI ^34^, coronary artery disease and type 2 diabetes^35^, and colorectal cancer^36^.

**Fig 2.**
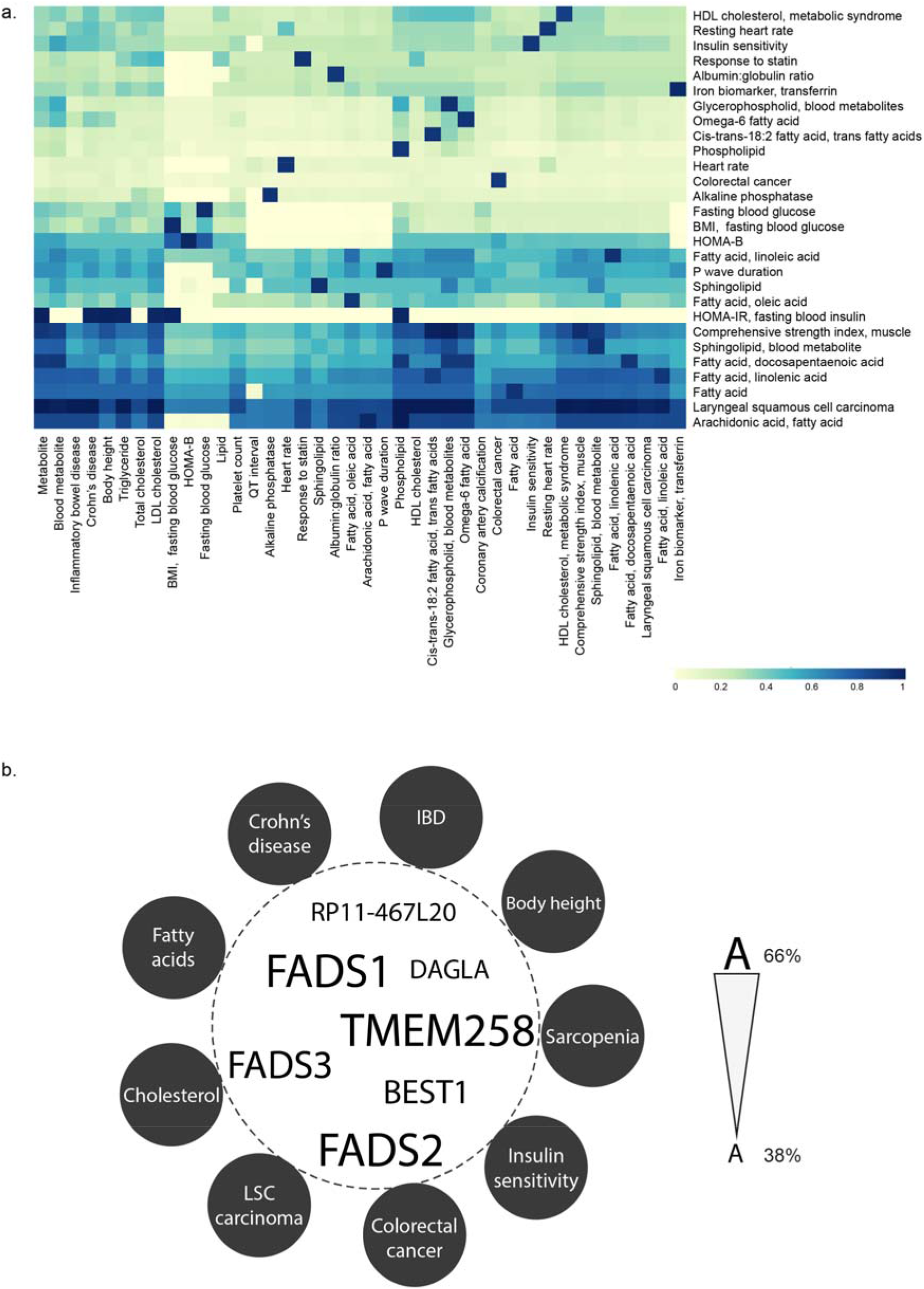
The largest eGene-defined multimorbidity cluster comprises forty phenotypes related to fat metabolism. (a) SNPs associated with inflammatory bowel disease, cholesterol levels, insulin sensitivity, muscle strength index, and laryngeal squamous cell carcinoma, amongst others, are associated with transcription effects at a pool of shared, multimorbid eGenes. Deep blue squares indicate higher proportions of shared eGenes, with 1 being the highest and indicating that two phenotypes have the same set of eGenes. (b) The *FADS1* locus on chromosome 11 is responsible for the identified inter-relationships *i.e*. the multimorbidity) between the phenotypes in this cluster. We hypothesize that the interrelationships between the additional 2,431 eGenes that are associated with these multimorbid phenotypes contribute to the unique characteristics and sub-clustering of the phenotypes (S3 Table). In this word cloud, the size of a gene name represent the percentage of phenotypes whose associated variants spatially affect the expression of that gene in the cluster e.g. the expression of *FADS1, FADS2* and *TMEM258* are associated with eQTLs in 26 of the 40 phenotypes in the cluster (S3 Table).

### Common genes are affected by different disease eQTLs

Variant rs174557 has been shown to moderate the binding of transcription factor *PATZ1* to downregulate of *FADS1*^37^. However, the effect of genetic variation on the regulatory network for the multimobid phenotypes associated with the *FADS1-3* locus has yet to be identified. Therefore, we mapped the eQTL-eGene interactions for which we had both spatial (*i.e*. Hi-C) and functional evidence (*i.e*. GTEx) within the FADS1-3 locus (Fig 3a). The transcription levels of the *FADS1, FADS2, TMEM258, and DAGLA* genes, that are central to this cluster, are associated with eQTLs that are located within these genes and across the region (Fig 3a). Nine of the putative regulatory regions are located within introns of genes (*i.e. MYRF, TMEM258, FEN1, FADS1, FADS2 and FADS3*) located within this locus. Putative regulatory effects linking eQTLs in *FADS1-3* with *DAGLA*, or eQTLs in *MYRF* with *FADS1-3* cross a topologically associating domain (TAD) boundary located in the vicinity of *FEN1*, whose transcription is not associated with any of the eQTLs (S8 Fig). eQTLs associated with some phenotypes (*e.g*. LDL cholesterol, muscle measurement and comprehensive strength index) are few and localised while others (*e.g*. cis-trans-18:2 fatty acid, phospholipid) are dispersed across the locus. However, despite this, almost all of the phenotype associated SNPs in this cluster are associated with a change in the transcript level of more than one gene (Fig 3b; compare with the pulmonary cluster, c in S6 Fig). Collectively, these results are consistent with previous reports of the formation of complex networks of multiple long-range interactions by regulatory elements and gene promoters^38^.

**Fig 3.**
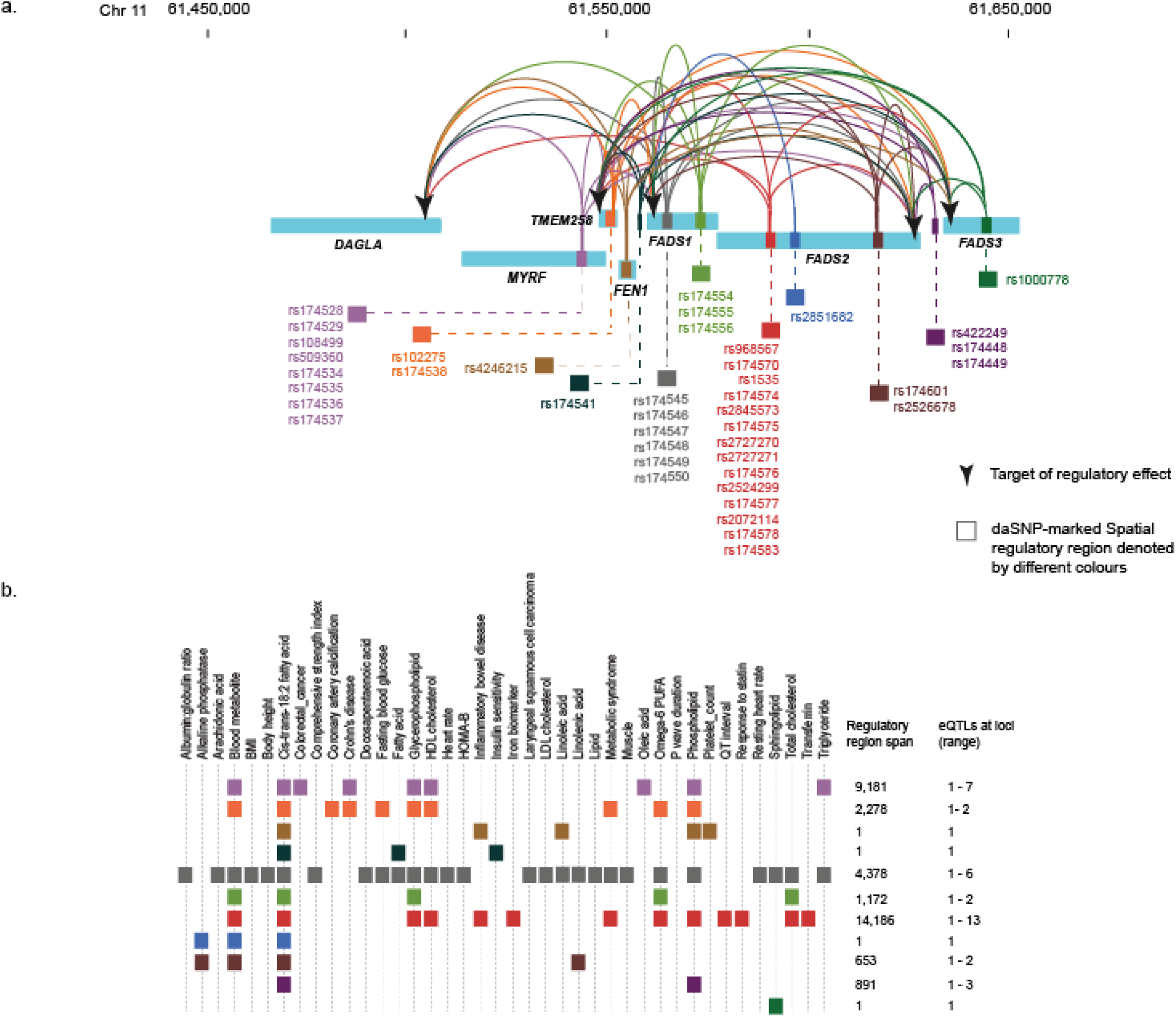
Spatial eQTL-eGene interactions central to the fat metabolism cluster. (a) Transcript levels for genes located within Chr 11: chr11:61447905 - 61659017 are associated with eQTLs located in clusters across the 283 kb locus. For simplicity, we grouped eQTLs according to separation in the linear sequence such that they are located in different genes, or are separated by ≤ 5 kb. Genomic locations are from human genome Hg19 and the eQTL-eGene interaction analysis used GTEx v4 (18/10/2016). (b) Phenotype-associated eQTLs are localized or dispersed across the *FADS1* locus, eQTLs associated with cis-trans-18:2 fatty acid levels are the most dispersed. eQTLs rs174545-50 are associated with the most phenotypes. eQTLs are coloured according to groups in A.

### eQTLs have gene and tissue specific effect patterns

SNPs that are in high linkage disequilibrium (LD) might be predicted to have inseparable regulatory effects on target genes. However, given the composite nature of regulatory elements and networks, it is likely that even linked SNPs affect different regulatory elements. Therefore, we obtained the effect sizes of eQTLs within the PUFA eGene cluster from different tissues (GTEx v7, 01/12/2017) to identify associations between eQTLs that were in strong LD. We characterised 12 distinct patterns of eQTL associations with the target genes within the *FADS* region (Fig 4a, S9 Fig). The eQTL effect patterns seem to be largely LD-dependent with some exceptions. For example, in the CEU population, rs174574 in Group E and rs1535 in Group L are in complete LD (*r*^2^ = 1) but have opposite effects on their common target genes. The minimum and maximum effect sizes of rs1535 are −0.76 (*FADS1, p*_eQTL_ = 5.3 × 10^−21^, cerebellum) and 0.8 (*FADS2, p*_eQTL_ = 2.3 × 10 ^−10^, spleen, while the maximum and minimum effect sizes of rs174574 are −0.72 (*FADS2, p*_eQTL_ = 1.5 × 10^−8’^spleen) and 0.73 (*FADS1, p*_eQTL_ = 1.5 − 10^−18^, cerebellum). By contrast, rs1000778 and rs422249 are not in LD (CEU *r*^2^ = 0.365) but have a similar effect pattern (Group B) on target genes, *FADS1* and *FADS2*. We observed similar patterns linking LD and regulatory effects within the *CHRNA* locus (a and b in S6 Fig).

**Fig 4.**
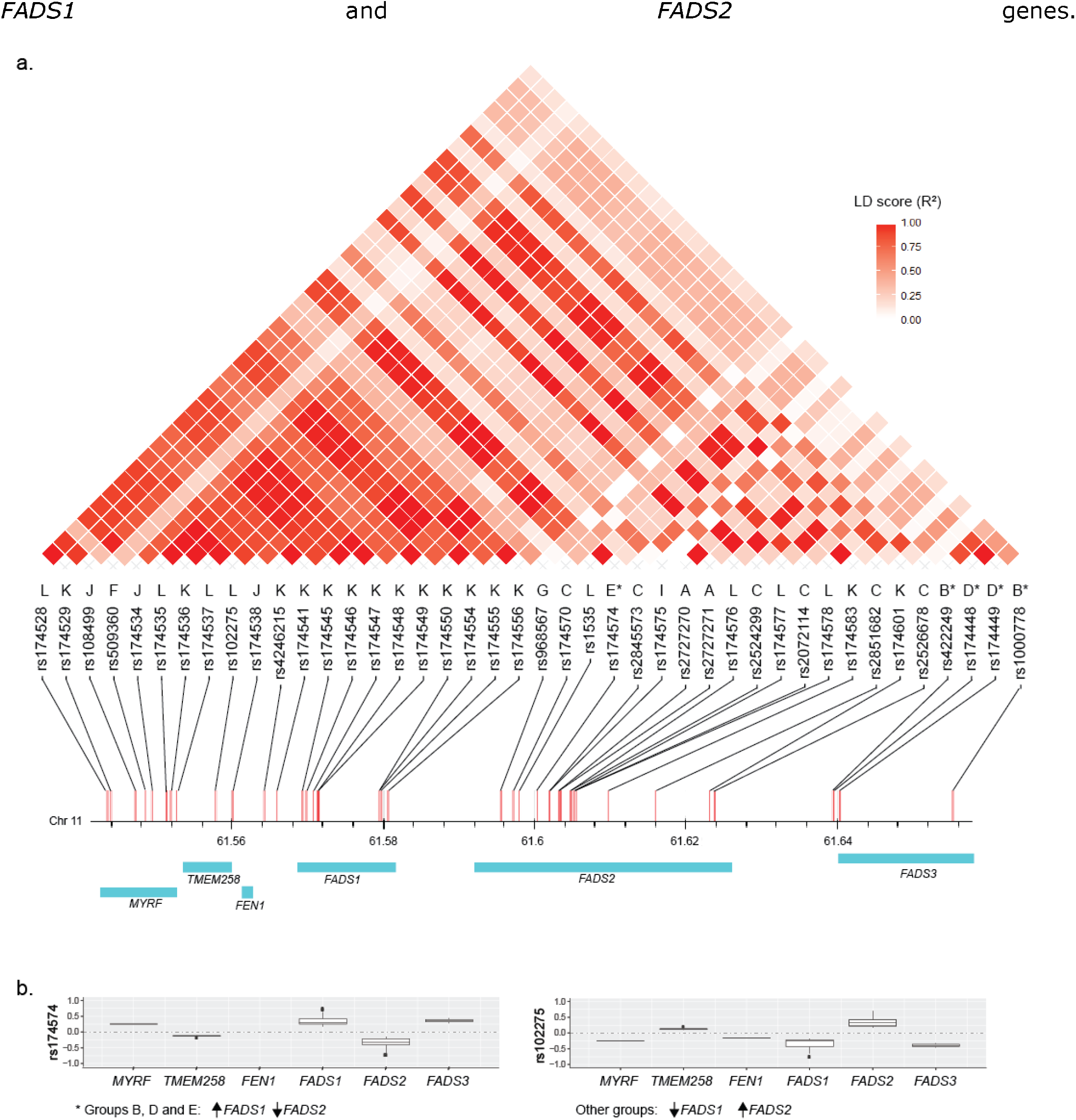
eQTLs have different effect patterns within the fat metabolism cluster. (a) There are 12 distinct eQTL effect patterns across the *FADS* locus. Categories A-L highlight the distinct pattern of eQTL associated transcriptional affects (S9 Fig). (b) There is an inverse association between the effects on *FADS1* and *FADS2* in all 12 eQTL effect patterns, B, D, and E patterns are associated with an increase in *FADS1* and decrease in *FADS2* transcript levels. All other eQTL patterns are associated with a decrease in *FADS1* and increase in *FADS2* transcript levels.. Effect sizes of spatial eQTL on eGenes were obtained from GTEx v7.

The fat metabolism cluster associated eQTLs located within the *FADS1-3* locus are all associated with an inverse relationship between *FADS1* and *FADS2* transcription levels. Notably, all but five eQTLs (*i.e*. rs174574, rs422249, rs174448, rs174449, and rs100078) are associated with a negative effect on *FADS1* and a positive effect on *FADS2* transcript levels (Fig 4b). Our results indicate that a composite regulatory hub forms from dispersed locations to regulate the convergent, duplicated (61% amino acid identity and have 75% similarity^32^)

Consistent with our understanding of tissue and cell type specificity regulation, the tissue eQTL effects showed cell line dependent enrichment patterns (Fig 5a; compare with a in S10 Fig). The strongest and weakest eQTL effects in subcutaneous adipose were observed in the GM12878 and KBM7 cell lines respectively, while in omental visceral adipose, they were observed in the HMEC and NHEK cell lines. The observed frequency of Hi-C contact counts of the eQTL-eGene pairs in the cell lines showed less tissue specificity (Fig 5b; b in S10 Fig). There was a strong positive correlation (*r* = 0.87) between the percentage of spatial eQTL-eGene interactions and the number of RNASeq GTEx samples in tissues (c in S10 Fig). Collectively, these results suggest that the frequency of spatial eQTL-eGene contact in cell lines are relatively uniform across tissues but the eQTL effects in the cell lines are tissue specific. This is consistent with previous results that show that transcription is not necessary for the formation and maintenance of a spatial connection^39,40^

**Fig 5.**
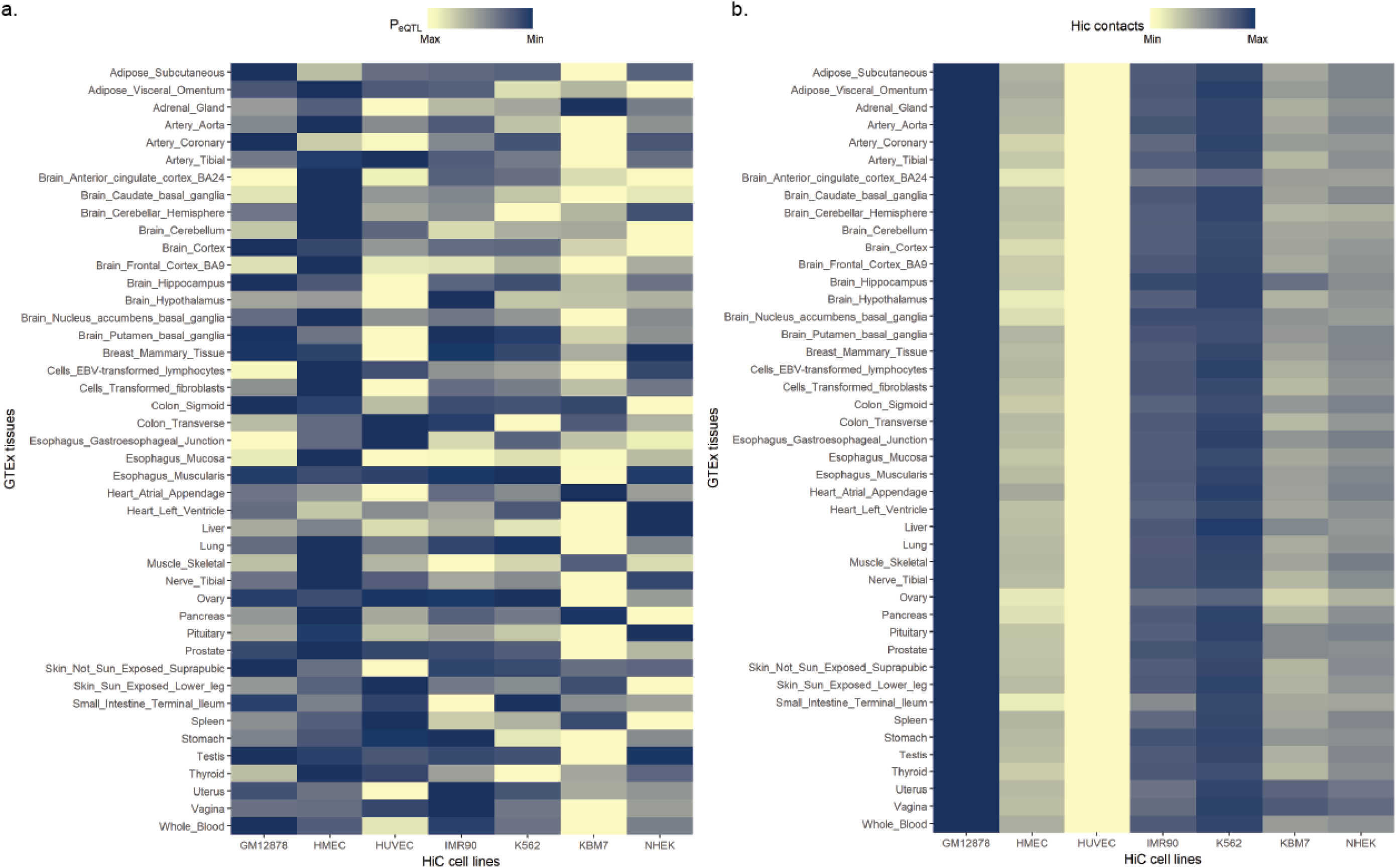
Tissue and cell line specific effects of eQTLs. (a) eQTL effects are strong in Hi-C cell lines that represent the tissues that they are derived from. The heatmap shows the range of mean eQTL p values in tissues, with the yellow and blue colours representing the HiC cell lines with the minimum and maximum mean p values respectively. (b) The number of HiC contact counts between the regions containing the eQTL-eGene pairs show less tissue specificity

### GWAS eQTLs spatially affect Mendelian genes

Rare monogenic or Mendelian diseases are typically considered to be associated with highly penetrant loss of function mutations. However, genes linked to Mendelian diseases have also been implicated in polygenic disorders^41–43^. Therefore, we determined if the spatial eGenes, we identified as being associated with complex diseases, were also implicated in Mendelian diseases. 62% (5,069) of the spatial eGenes we identified are catalogued in the OMIM database (Fig 6a, S4 Table). This is consistent with the possibility that there is a distal regulatory component in rare Mendelian disorders.

The gene-phenotype linkage mapping category in OMIM includes those genes whose exact locations and strands are yet to be resolved in major resources like the UCSC genome browser and Gene Cards. Our method captured a significantly low proportion (0.6%) of gene-phenotype associations that are mapped through linkage in OMIM (15.4% of the total gene-phenotype associations). By contrast, factoring in the 3D genome organization significantly increased the chance of identifying gene-phenotype associations (*i.e*. 81.9% to 98.2%) for which there is a known molecular basis (Fig 6b).

**Fig 6.**
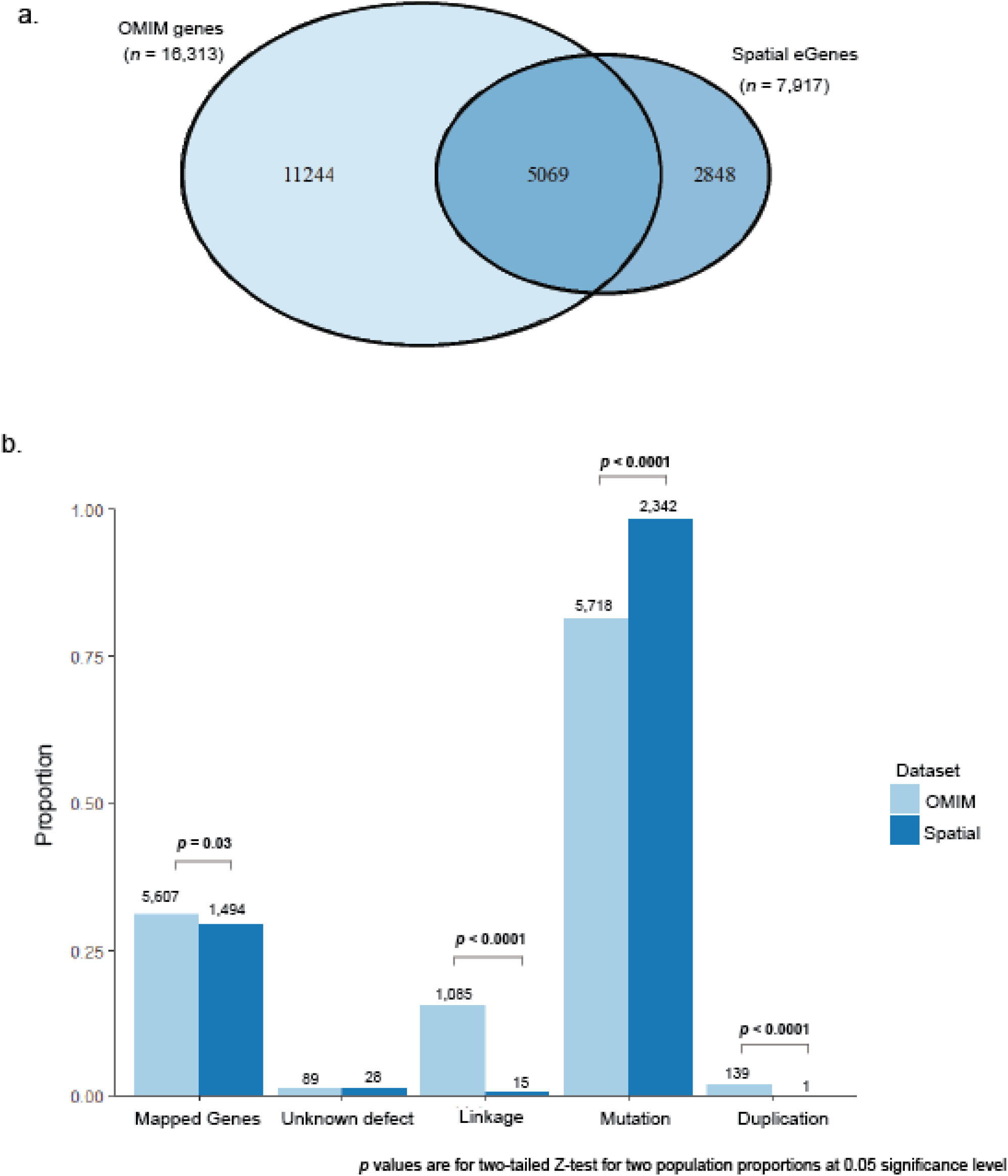
OMIM analysis of spatial eQTL-eGene interactions reveal differences in gene-phenotype mappings. (a) 62% of the spatial eGenes we identified are associated with human disease in the OMIM database (retrieved, 11/08/2017, S4 Table). (b) 98,2% of gene-phenotype associations of the identified spatial eGenes are based on known mutations in the genes.. OMIM mapping methods are: 1) Gene has unknown underlying defect but is associated with the disorder; 2) Disorder is mapped to gene based on linkage but mutation in gene has not been found; 3) A mutation in the gene has been identified as the basis of the mapped disorder; and 4) Disorder is caused by deletion or duplication of contiguous genes. Mapped gene proportions were calculated as the number of genes mapped to at least one phenotype in the OMIM data (annotated in graph) divided by the total number of genes in the OMIM data (16,313 for OMIM genes) or the number of spatial eGenes annotated in the OMIM data (5,069 for eGenes).

## Discussion

Here we created a genetic “multimorbidity atlas” of traits that have the same set of genetic components (*i.e*. eQTLs or eGenes) with no particular bias to causation, confounding or endpoint effects.

We identified greater pleiotropy in human complex diseases and phenotypes at the gene level than at variant level. Previous studies have reported pleiotropy in complex traits^44–46^. Our findings are consistent with the work of Sivakumaran et al^46^ in 2011, who reported 16.9% and 4.6% pleiotropy at the gene and variant levels. However, our study differs from theirs in significant ways as Sivakumaren *et al*.: 1) =used 1,687 SNPs that satisfied the GWAS significance threshold (p values < 5 × 10^−8^); 2) adopted the target genes that were suggested by the authors of the GWAS, annotated in the GWAS Catalog, or are in LD with tag SNPs; and 3) reported that variant pleiotropy is associated with gene location, and that exonic variants are more pleiotropic than intergenic variants. By contrast, we used 7,776 SNPs with suggestive GWAS p values (< 5 × 10^−6^) and defined the target genes using spatial eQTL evidence. Notably, we find that spatial eQTLs within 1 megabase of eGenes are more than twice as common as eQTLs within genes (S2 Fig). The integration of genomic organization information into the interpretation of SNP function enabled the identification of novel regulatory interactions in complex traits. Further empirical studies are required to validate these interactions.

We hypothesise that the gene pleiotropy we identified within the phenotype clusters drives the multimorbidity between the phenotypes. The most common genes in the multimorbid clusters are typically located adjacent to each other in a contiguous genomic region (Fig 4 and a in S6 Fig). We conclude that these regions are super regulatory loci that are comprised of different composite regulatory elements, each having a distinct and distinguishable effect on the genes therein. These effects are reflected in the LD architecture of the regions and indicates that the inheritance off these regions may be linked to physical association between regions that are separated in the linear sequence. Moreover, the finding of large effect sizes for eQTLs involving variants in genomic regions with low LD is consistent with previous observations of greater deleterious effects, and larger per-SNP heritability^47^ for poorly linked variants, while genomic regions with high LD have lower heritability and greater exonic deleterious effects^48^.

The concentration of multiple intronic eQTLs within low recombination cluster regions indicates inherited allelic heterogeneity (*i.e*. multiple signals at a locus that affect a trait)^49,50^. This is consistent with evidence that discrete multiple variants (and not a single causal variant) within an LD block impact multiple linearly separated enhancers and the expression of target genes ^11,51,52^. However, causative variants cannot be separated from disease modifiers at this level because LD is subject to allele frequency, recombination, selection, genetic drift and mutation^53,54^. As such, variants in LD can affect each other’s statistical values^55^.

Our finding that *FADS1* and *FADS2* are inversely associated with eQTLs located across the *FADS* locus informs on the mechanism through which genetic variation contributes to the biochemistry of PUFA synthesis in complex multimorbid disorders. *FADS1* and *FADS2* encode the delta-5 (D5D) and delta-6 desaturase (D6D) enzymes, respectively, which catalyse the rate-limiting steps in PUFA biosynthesis respectively^32,56^. Inhibition of D6D, which acts upstream of D5D in the pathway, has been correlated with decreases in inflammation in several rodent studies^56–58^. Our results are consistent with observations that an increase in copy number of the *FADS1* variant rs174548 (Group K; Fig 4; S9 Fig) correlates with: a) reduced efficiency of D5D; b) increased concentrations of phospholipids with PUFA sidechains having 3 or less double bonds^59^. Our results are consistent with a significant genetic contribution to reduced expression of D5D, which in turn leads to a build-up of pro-inflammatory eicosanoids (via n-6 PUFA). Notably, only 5 genetic variants (rs422249, rs174448, rs174449, rs1000778 and rs174574) increase D5D transcript levels and thus favour the synthesis of anti-inflammatory eicosanoids.

TAD boundaries are generally considered to be conserved across tissues and developmental stages^60,61^. However, differences in TAD formation do occur^62,63^. We observed both intra- and inter-TAD eQTL-eGene interactions, in addition to eQTLs involving variants located at TAD boundaries. For example, eQTLs rs8042374 in the *CHRNA* locus (about which the lung disorders cluster, S7 Fig) and rs174537 in the *FADS* locus (S8 Fig) both lie at a TAD boundary. This is consistent with observations that genetic mutations at TAD boundaries can impact on enhancer-promoter interactions^64,65^. It remains possible that cell line, developmental or cellular state specific chromatin interactions^66,67^ have been missed in the HiC libraries we used to identify the eQTL-eGene and inter-phenotype relationships. Future work should focus on tissue specific Hi-C library formation to enable the teasing apart of the nuances associated with cell and tissue specific chromatin interactions in complex disorders.

Several studies have shown that both large-effect rare variants and small-effect variants are associated with complex diseases^52,68,69^. Yet, there is no evidence that the rare variants located at the gene locus are the main drivers of the genetic variance^70^. Our analysis of the OMIM database suggests that genes harbouring rare variants with large effects are also distally regulated by common variants with small effects.

We hypothesise that the common genes within a phenotype cluster highlight the underlying molecular mechanisms that drive shared multimorbidity (Fig 7a). By contrast, the unique presentations of individual phenotypes within a multimorbid cluster result from molecular mechanisms driven by genes that are not shared by other members of the cluster (Fig 7a). The genetic contribution to the regulation of multimorbidity is explained by three nonexclusive models: 1) genetic variants that are associated with more than one disease phenotype affect the same target genes (Fig 7b), indicating genetic pleiotropy; 2) different genetic variants associated with the multimorbid phenotypes mark a single regulatory element (*e.g*. a super-enhancer) and thus common gene(s) (Fig 7c); or 3) different variants each marking different regulatory elements that target the same gene (Fig 7d).

**Fig 7.**
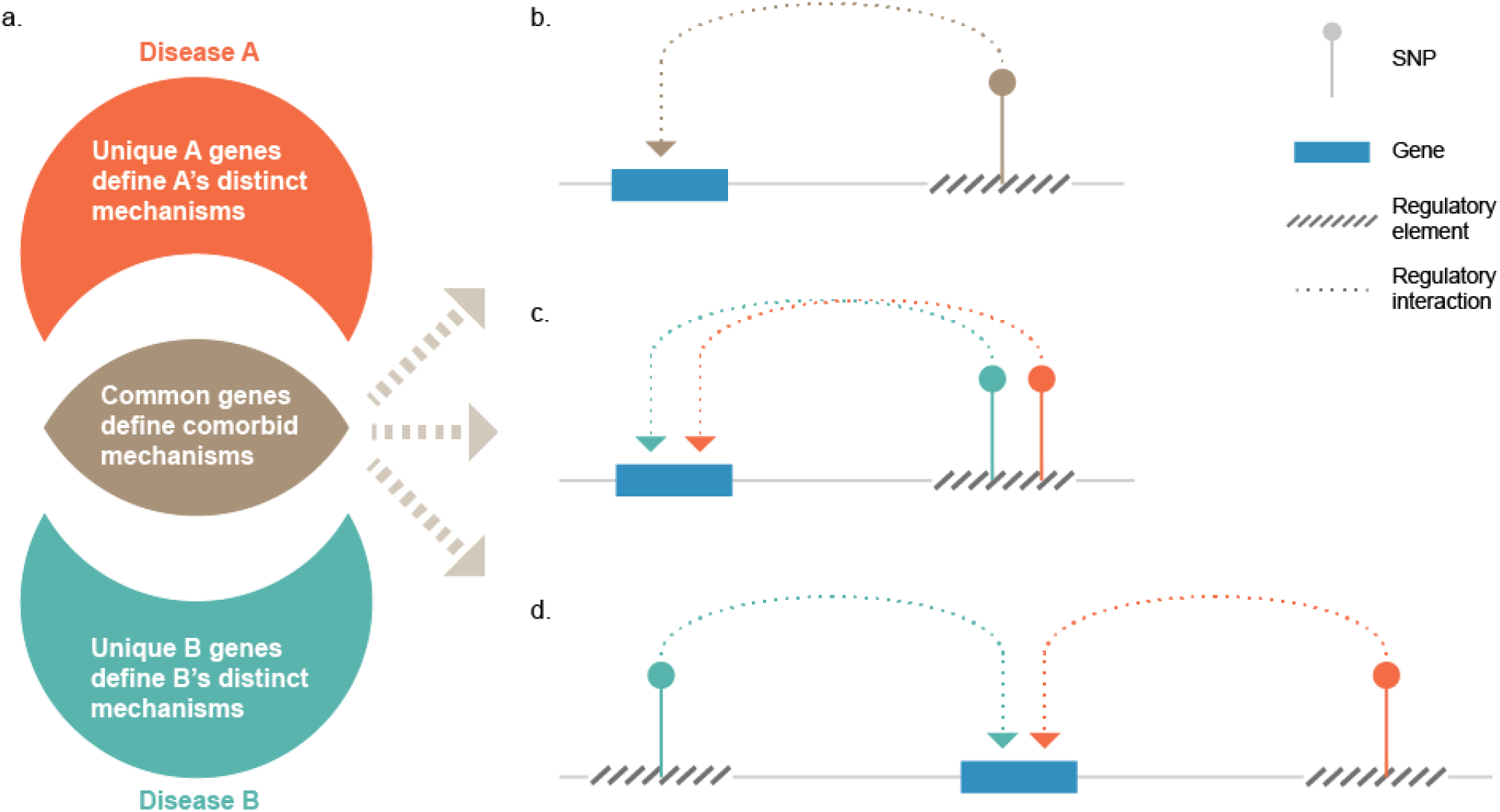
Schematic model of gene pleiotropy in multimorbidities. (a) Common target genes between any two complex disorders highlight the molecular mechanism(s) that underlie their common pathogenicity. The sets of target genes that are unique to the disorders represent the mechanisms that make the disorders different. Gene pleiotropy in multimorbidities of complex disorders can occur when (b) a variant associated with the disorders marks a regulatory element that target the common gene; (c) different variants associated with different disorders mark the same (super)-regulatory element that impacts the common gene; or (d) different regulatory elements marked by different variants impact the common gene.

In conclusion, the integration of spatial and gene eQTL information with phenotype association data leads to the identification of the genetic components that encode the molecular mechanisms that underlie both the multimorbidity and the unique development of complex disorders and traits. Further refinement of these relationships will require empirical studies that integrate multi-omics and epigenetic information on cells and tissues from patients with multimorbid disorders.

## Methods

### Identification of spatial eQTL-eGene pairs

All genetic variants (*p* value < 5 ×10^−6^) from GWAS associations (www.ebi.ac.uk/qwas; v1.0.1, downloaded on 25/08/2016) were run through the CoDeS3D pipeline as previously described (Fadason et al., 2017), for the identification of eGenes that are affected by spatial eQTLs and the tissues in which those interactions occur. In summary, high resolution HiC chromatin interaction libraries of GM12878, HMEC, HUVEC, IMR90, K562, KBM7 and NHEK lines ^21^ were interrogated for genes in chromatin regions that physically interact with SNP regions. The resulting SNP-gene pairs were used to query the GTEx database (www.atexportal.org. multi-tissue eQTLs analysis v4) to identify eQTL-eGene pairs i.e. SNPs that are associated with a change in the expression genes.

### Construction of phenotype matrices

A mXn matrix of a_*i,j*_ was constructed, where m and n are the same set of phenotypes and a is the proportion of eGenes in phenotype *i* that are common with phenotype *j*. We defined a phenotype as the trait associated with a SNP in the GWAS Catalog. Sometimes more than one trait are associated with a SNP in a single study, in that case we created a composite phenotype of the traits. The pairwise ratio of common eGenes between phenotype *i* and phenotype *j* was calculated as the number of their common genes divided by the total number of eGenes associated with phenotype *i*,

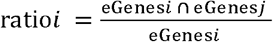

A similar matrix of pairwise eQTL ratios was also constructed.

To control for the eGene matrix, all 7,917 eGenes were pooled together and randomly assigned to phenotypes so that each phenotype in the control matrix had the same number of eGenes as its corresponding phenotype in the eGene matrix. The pairwise ratios of common eGenes among the phenotypes were calculated as done in the eGene matrix. 1000 different null datasets were constructed in this manner and the mean matrix was calculated.

### Convex biclustering of phenotypes

To group phenotypes based on the eGenes they share, we selected only phenotypes that have ≥ 4 eGenes in common. We used the cvxbuclustr R package, a convex biclustering algorithm that simultaneously groups observations and features of high-dimensional data ^24^. A combined Gaussian kernel with k-nearest neighbour weights of the phenotype eGene ratios matrix was constructed. A biclustering solution path of 100 equally spaced y parameters from 10^0^ to 10^3^ was initialised and a validation using the `cobra_validate` function was performed to select a regularization parameter γ, on which the biclustering models would be based. The `biclust_smooth` function was used to generate a bicluster heatmap of data smoothed at the model with the minimum validation error (Uγ^⋆^)

### Multimorbidity Analysis

To identify eGenes that are central to phenotype biclusters, an eGene commonality index was calculated for each eGene in the cluster. We defined the commonality index of an eGene as the ratio of phenotypes in a cluster that are associated with that eGene. Mapping of eQTL-eGene interactions and their effects in the fat metabolism cluster was based on GTEX v7 multi-tissue analysis and hg19 genome assembly respectively. The linkage disequilibrium analysis of eQTLs in the FADS region was done on CEU population data obtained from LDLink 3.0 (https://analysistools.nci.nih.gov/LDIink/)^71^. Visualization of HiC, H3K4me1, H3K27ac, DNAse and Pol2 data in the GM12878 cell line was done with HiGlass (higlass.io)^72^.

### OMIM Analysis

The ‘genemap2’ data was obtained from the OMIM database (omim.org, accessed 11/08/2017). Spatial eGenes that are included in the OMIM database were analysed for gene-phenotype mapping methods and compared with the OMIM genes. OMIM’s phenotype-gene mapping methods are numbered thus: 1) Gene has unknown underlying defect but is associated with the disorder. 2) Disorder is mapped to gene based on linkage but mutation in gene has not been found. 3) A mutation in the gene has been identified as the basis of the mapped disorder. 4) Disorder is caused by deletion or duplication of contiguous genes.

### URLs

GWAS Catalog: https://www.ebi.ac.uk/awas/

GTEx portal: https://www.gtexportal.org/home/

LDLink 3.0: https://analvsistools.nci.nih.gov/LDIink/

HiGlass: higlass.io

HiC data^21^ GEO accession number: GSE63525.

**Life Sciences Reporting Summary**

### Data and Code Availability

CoDeS3D pipeline is available at https://github.com/alcamerone/codes3d.

Supplementary tables are available at https://fiqshare.com/s/4d53e690d9adde385057

Scripts used for data curation, analysis, and visualisation are available at https://github.com/Genome3d/multimorbidity-atlas.git

## Acknowledgements

This work was funded by High Value Nutrition National Science (MBIE/HVN grant #3710040) to JOS & TF. JOS and WS are funded by a Royal Society of New Zealand Marsden Fund [Grant 16-UOO-072],

## Author contributions

TF ran analyses, wrote software, interpreted data and wrote the manuscript. WS contributed to data interpretation and commented on the manuscript, JOS directed the study, contributed to data interpretation and co-wrote the manuscript. JOS is guarantor for this article.

## Supporting Information

**S1 Fig.**
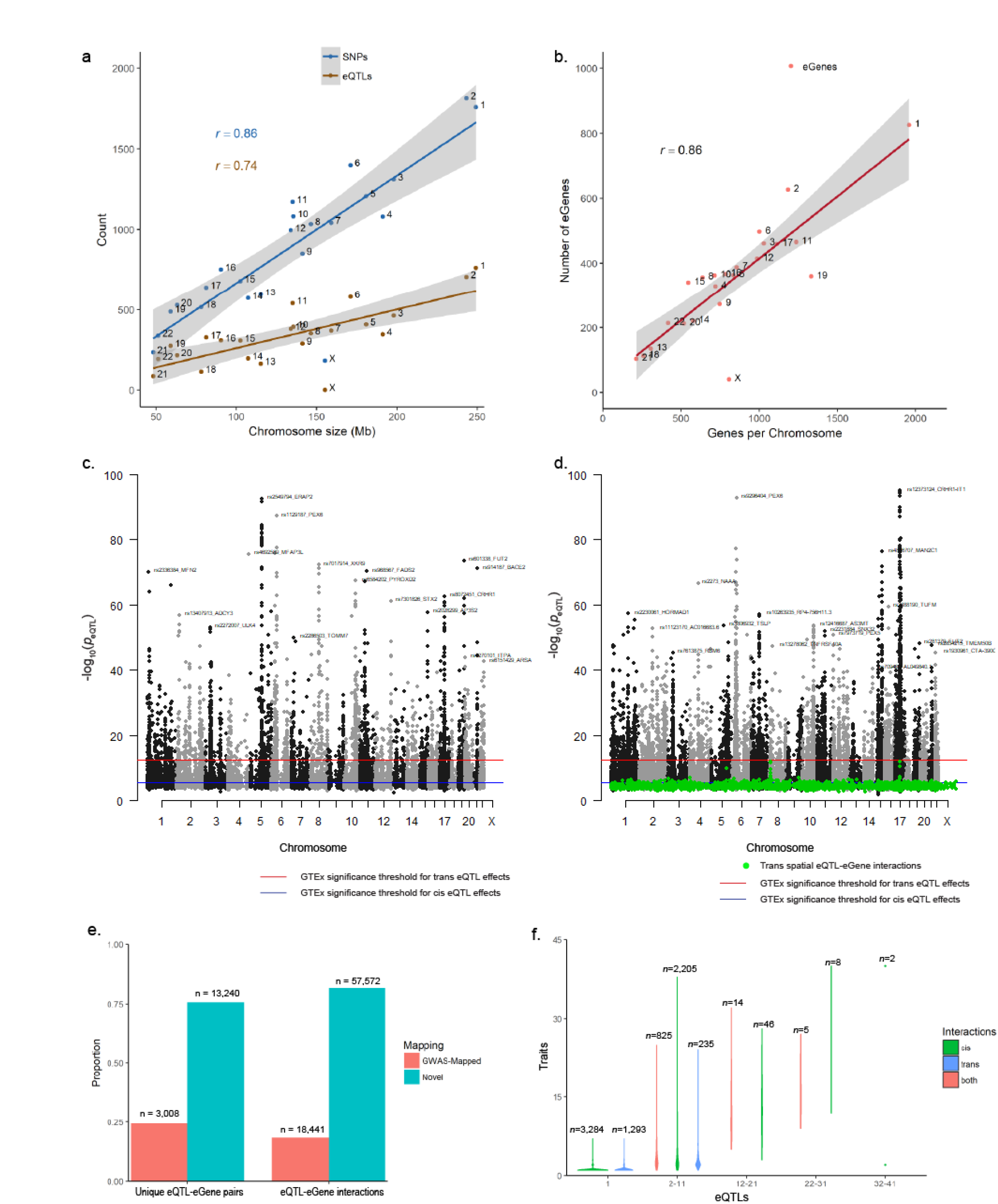
Summary of eQTL-eGene interactions. (a) Correlation between chromosome size with number of SNPs (*n* = 20,782) and eQTLs (*n* = 7,776). (b) Correlation between number of eGenes on chromosomes and the total number of genes on chromosomes. (c) Manhattan plot showing the eQTL *p* values (GTEx v4) of significant (*FDR* ≤ 0,05) GWAS-mapped eQTL-eGene interactions in tissues. (d) Manhattan plot of significant eQTL-eGene interactions that are not mapped in the GWAS Catalog, We considered an interaction as novel if the eGene is not distinctly mentioned in the ‘MAPPED_GENE’ column of a SNP association in the GWAS Catalog (v1.0,1). (e) Proportions of the GWAS-mapped and novel eQTL-eGene pairs and their interactions in GTEx tissues. (f) Violin plots of eGenes highlighting the number their associating phenotypes and the number of eQTLs affecting them. Interactions are as *cis*-only (*i.e*. eQTL and eGene are ≤ 1Mb apart), *trans*-only (*i.e*. eQTL and eGene are > 1Mb apart or on different chromosomes) or both.

**S2 Fig.**
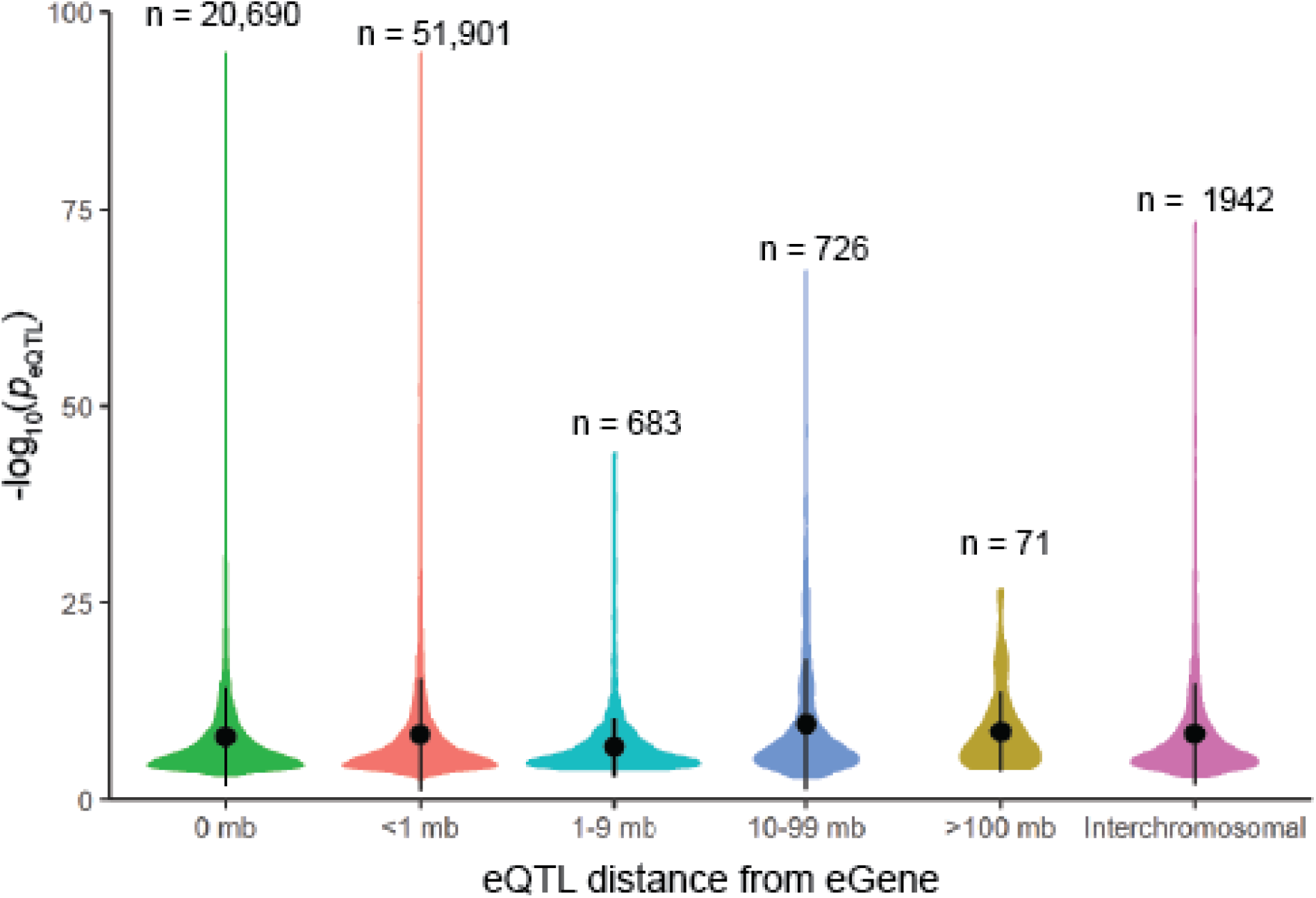
Violin plot of eQTL p values and distance between eQTL and eGene interactions. There are more eQTLs within 1 mb of genes than there are within genes.

**S3 Fig.**
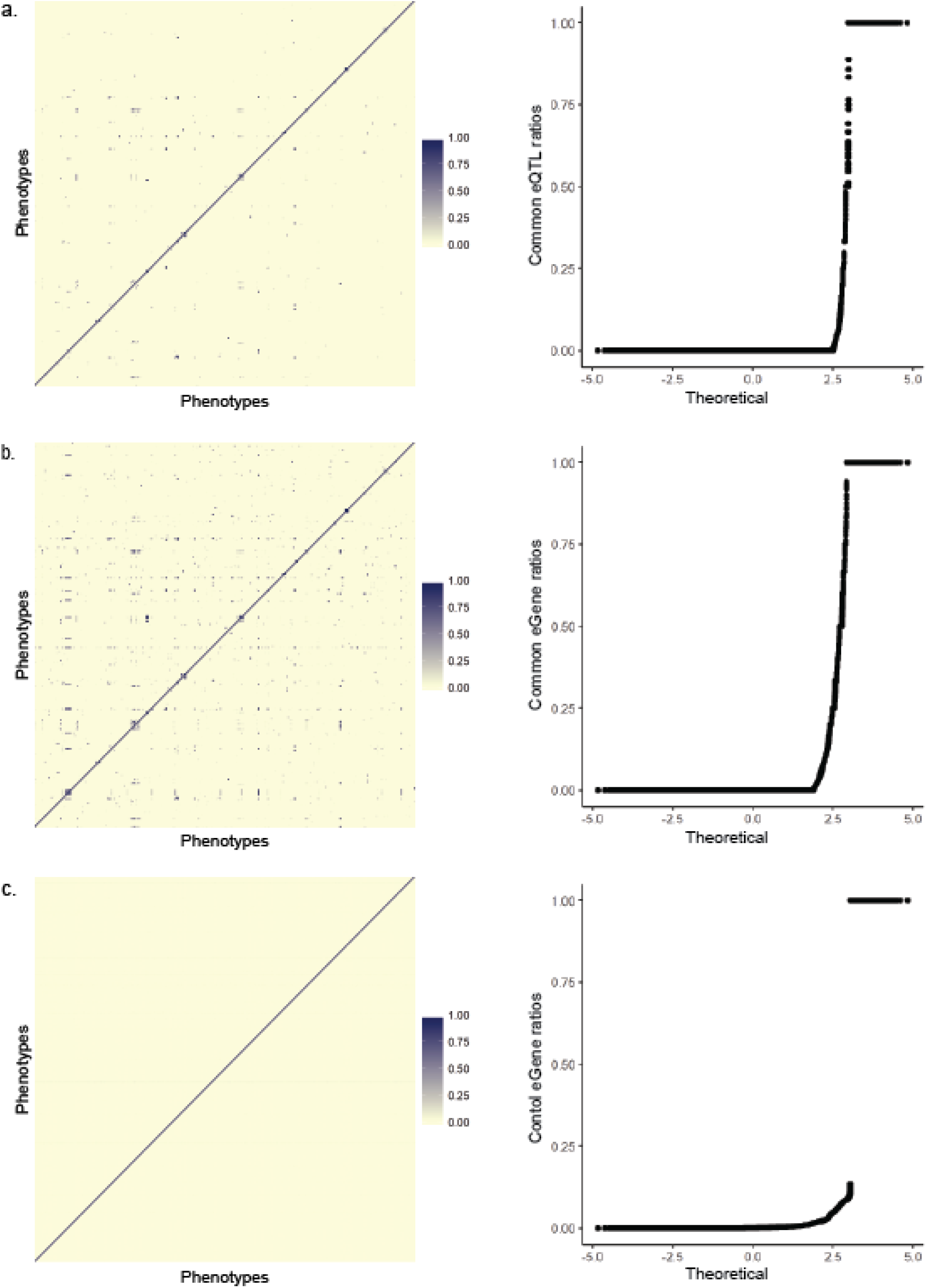
Associations of phenotypes based on shared spatial interactions. (a) Phenotypes associate weakly when the 7,776 significant eQTLs were used to define their interrelationship as shown by sparser blue dots in the heatmap and gaps on the Q-Q plot. (b) Relationships among phenotypes are enhanced by the eGenes. (c) 1000 null datasets were generated by randomly assigning eGenes to phenotypes such that each control phenotype has the same number as the corresponding sample phenotype. The mean null dataset has a different pattern from that of sample phenotypes. Heatmaps in a, b and c are extracts of the same set of phenotypes in the same order. Darker squares in matrices indicate higher proportions of shared eGenes (with 1 being the highest, meaning the sets of eGenes of two phenotypes are the same). Q-Q plots include shared eQTL, eGene or control ratios for 861 phenotypes.

**S4 Fig.**
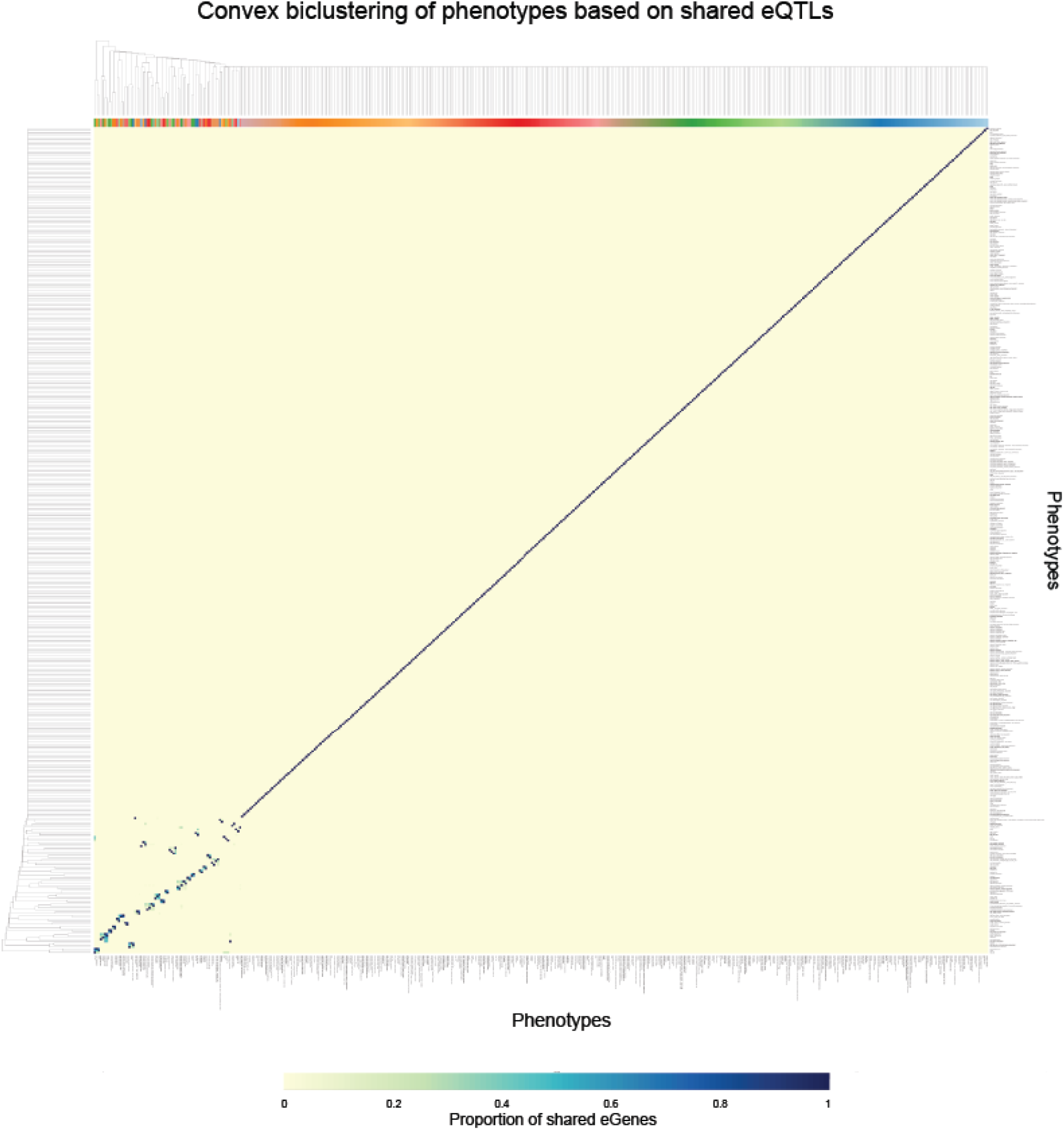
Biclustering of phenotypes cluster based on the shared eQTLs. The graph shows the segregation of phenotypes (on both axes) that share ≥ 4 eQTLs with at least one other phenotype by the convex biclustering algorithm from the cvxbiclustr R package. Deep blue squares indicate higher proportions of shared eGenes, with 1 being the highest and indicating that two phenotypes have the same set of eGenes. The complete proportions of the eGenes shared among phenotypes are given in S2 Table.

**S5 Fig.**
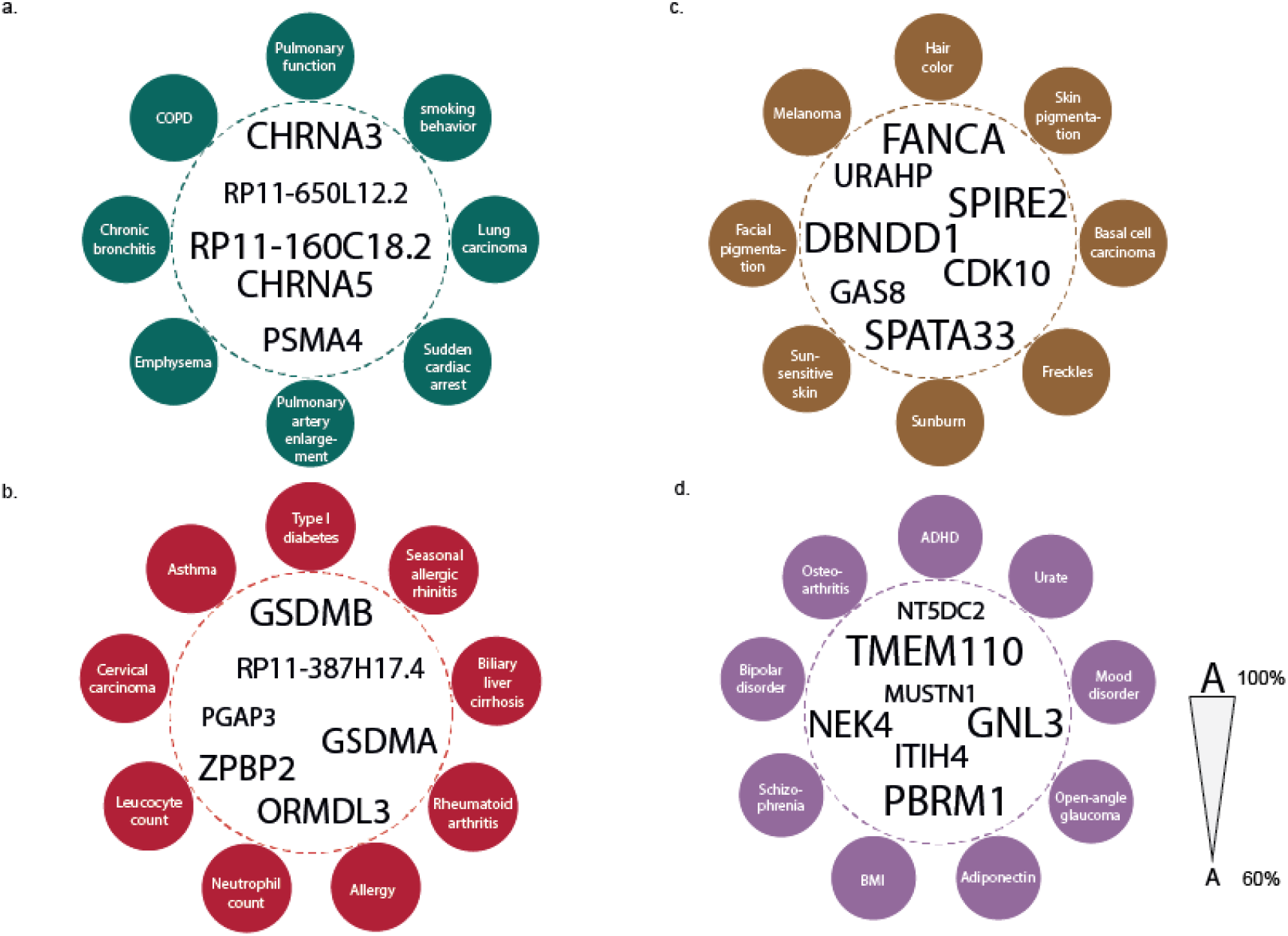
Examples of phenotype clusters. (a) The *CHRNA5 – ADAMTS7* region (chromosome 3:78,832,747 – 79,103,773; hg19) is important for the clustering of disorders of pulmonary function: chronic obstructive pulmonary disease, smoking behaviour, lung carcinoma and sudden cardiac arrest. (b) Immune disorders such as type 1 diabetes, cervical cancer, rheumatoid arthritis, and biliary liver cirrhosis are clustered around the *PGAP3 – GSDMA* region on chromosome 17: 37,827,375 – 38,134,431. (c) Skin pigmentation, hair colour, facial pigmentation, melanoma, basal cell carcinoma, freckles and sunburn share genes in the *SPATA33 – URAHP* (chromosome 16:89724152 – 90114191) region. (d) Genes from the *NT5DC2 – TMEM110* (chromosome 3:52558385 – 52931597) region are common to mental disorders, body mass index, osteoarthritis, open-angle glaucoma and adiponectin measurement. Proportions of eGenes for the four clusters are given in S3 Table.

**S6 Fig.**
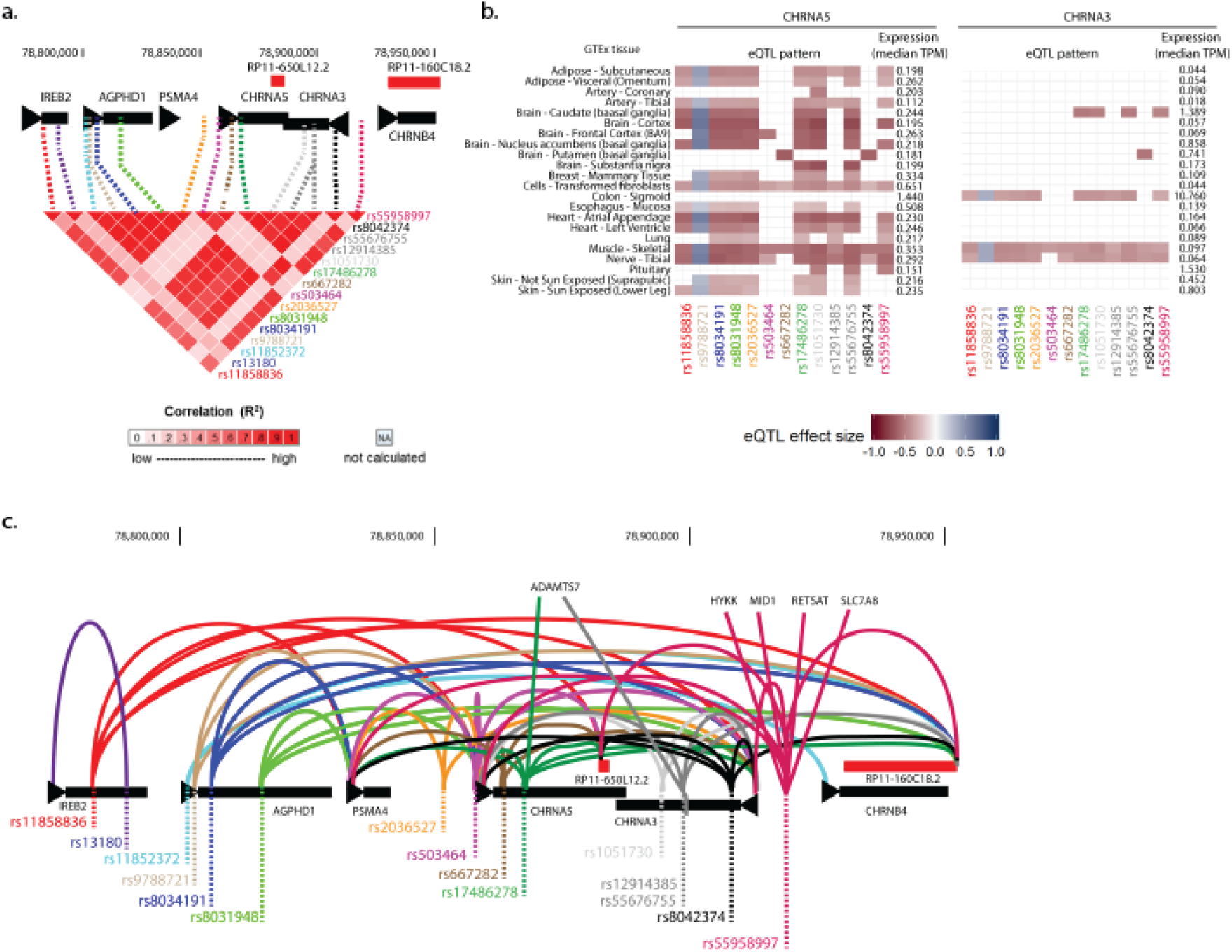
The CHRNA regulome is central to pulmonary disorders. (a) The LD pattern here suggests 2 alternating haplotype blocks, with the smaller block represented by rs503464, rs667282, rs8042374, and to a lesser extent, rs13180. (b) CHRNA3 and CHRNA5 are differentially expressed and regulated in the same tissues and by the same eQTLs, which seem consistent with the LD patterns in a. (c) eQTLs in the CHRNA locus region interact with multiple genes both within and outside this region, suggesting that the locus may be a super-enhancer.

**S7 Fig.**
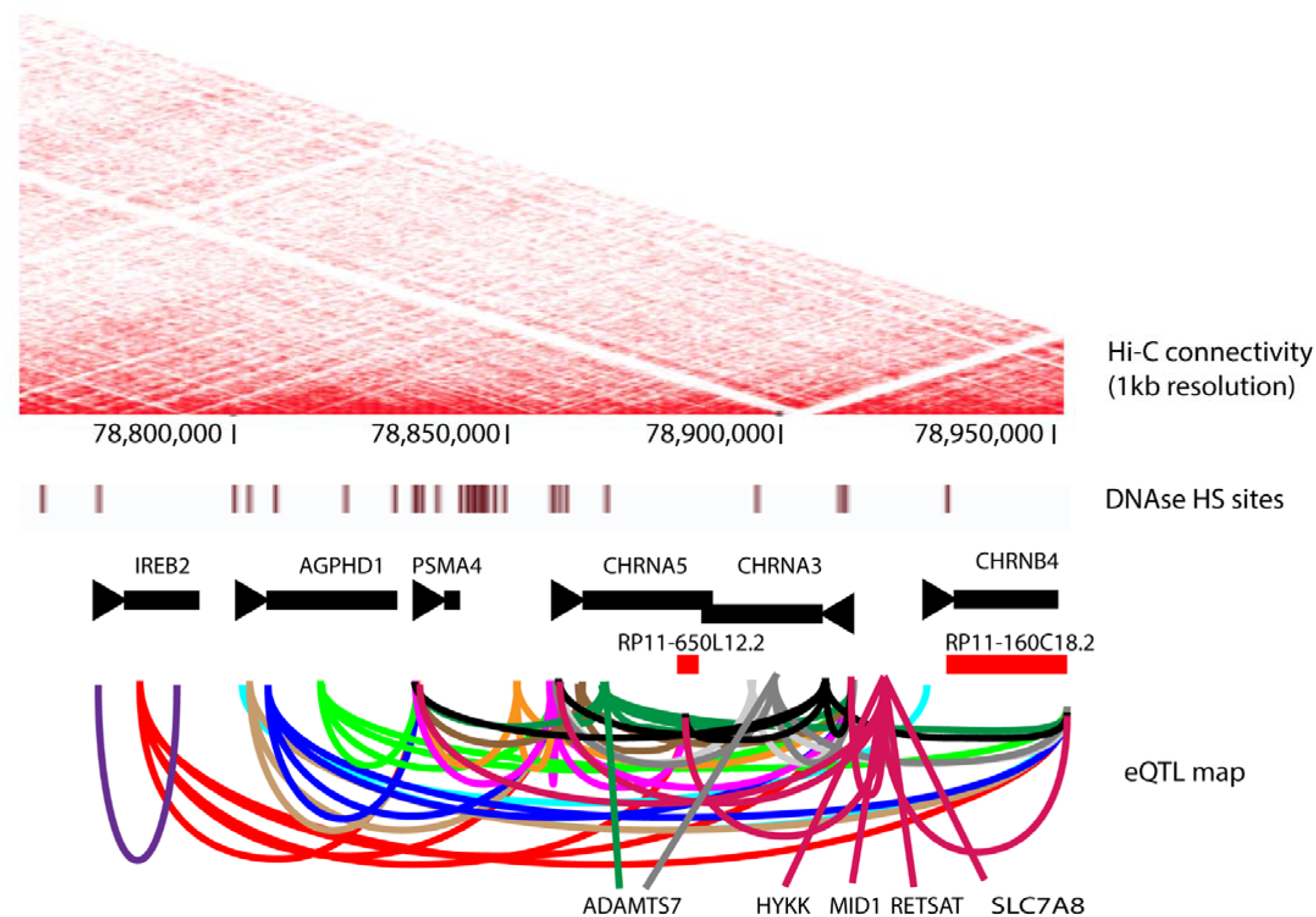
eQTLs in the CHRNA locus are involved in intra- and inter-TAD interactions. rs8042374, which is located at a TAD boundary also interacts with genes in the two TADs.

**S8 Fig.**
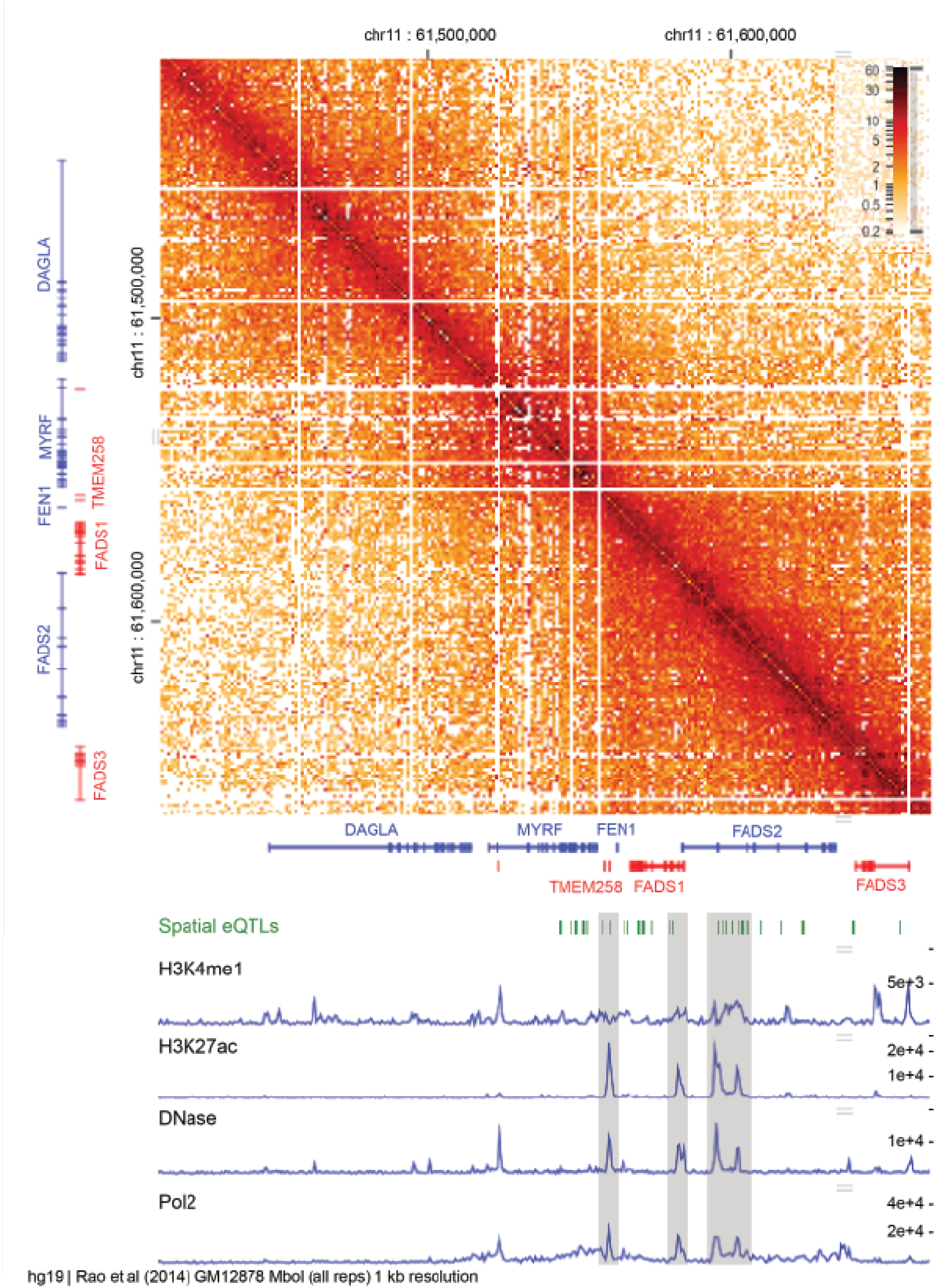
eGenes are affected distally by eQTLs. (a) eQTLs in the fat metabolism cluster are found in putative enhancer regions and have inter-TAD effects on genes, HiC image is was produced with HiGlass (higlass.io)

**S9 Fig.**
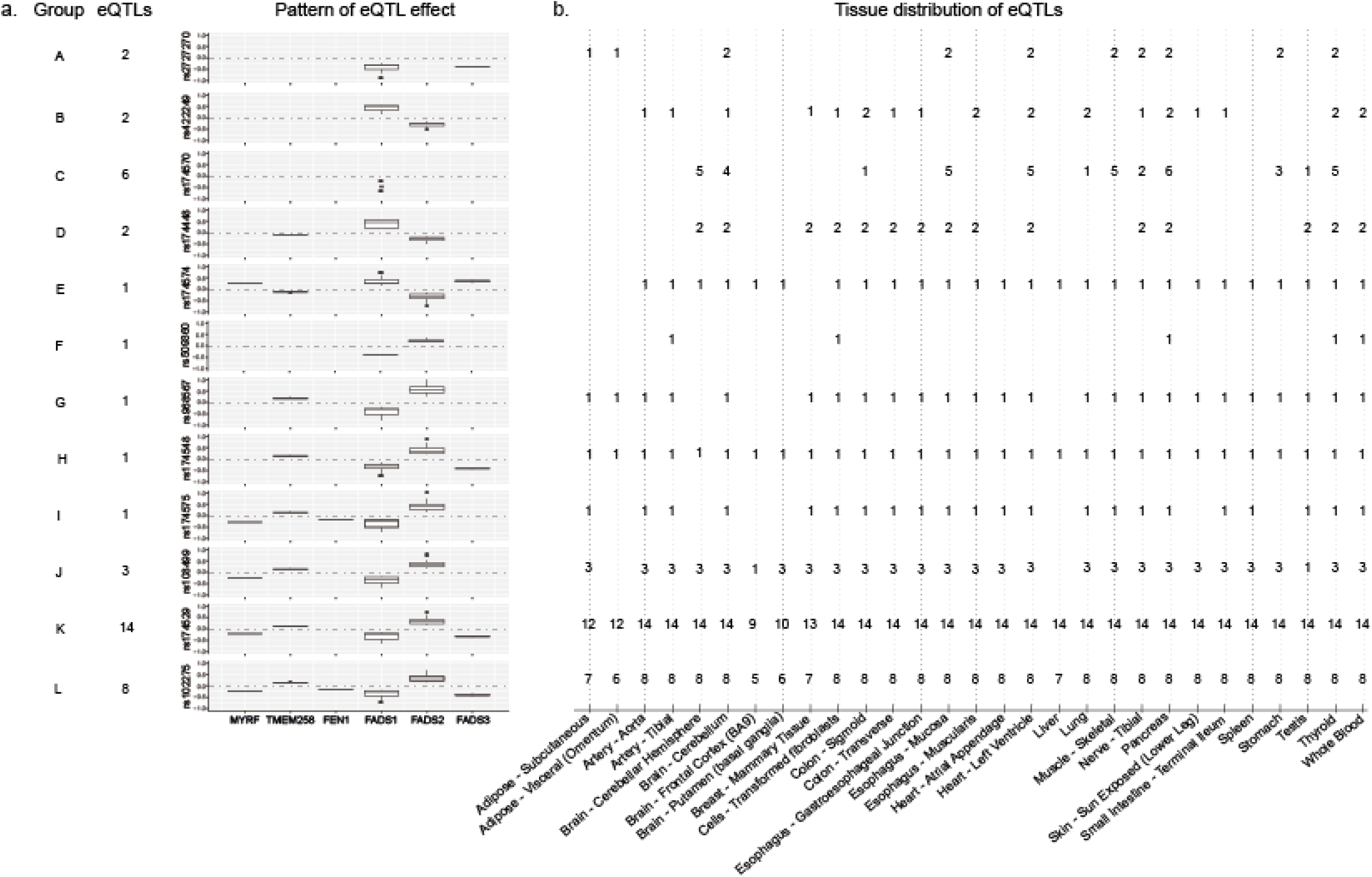
eQTLs have different patterns of effect on genes in the fat metabolism cluster. (a) Box plots of the effect sizes of eQTLs on genes revealed 13 different patterns as shown by a representative eQTL for each group (See S4 Table for grouping of eQTLs). (b) The tissue distribution of the eQTLs seem to differ based on their effect size patterns. Effect sizes of spatial eQTL on eGenes were obtained from GTEx v7 analysis.

**S10 Fig.**
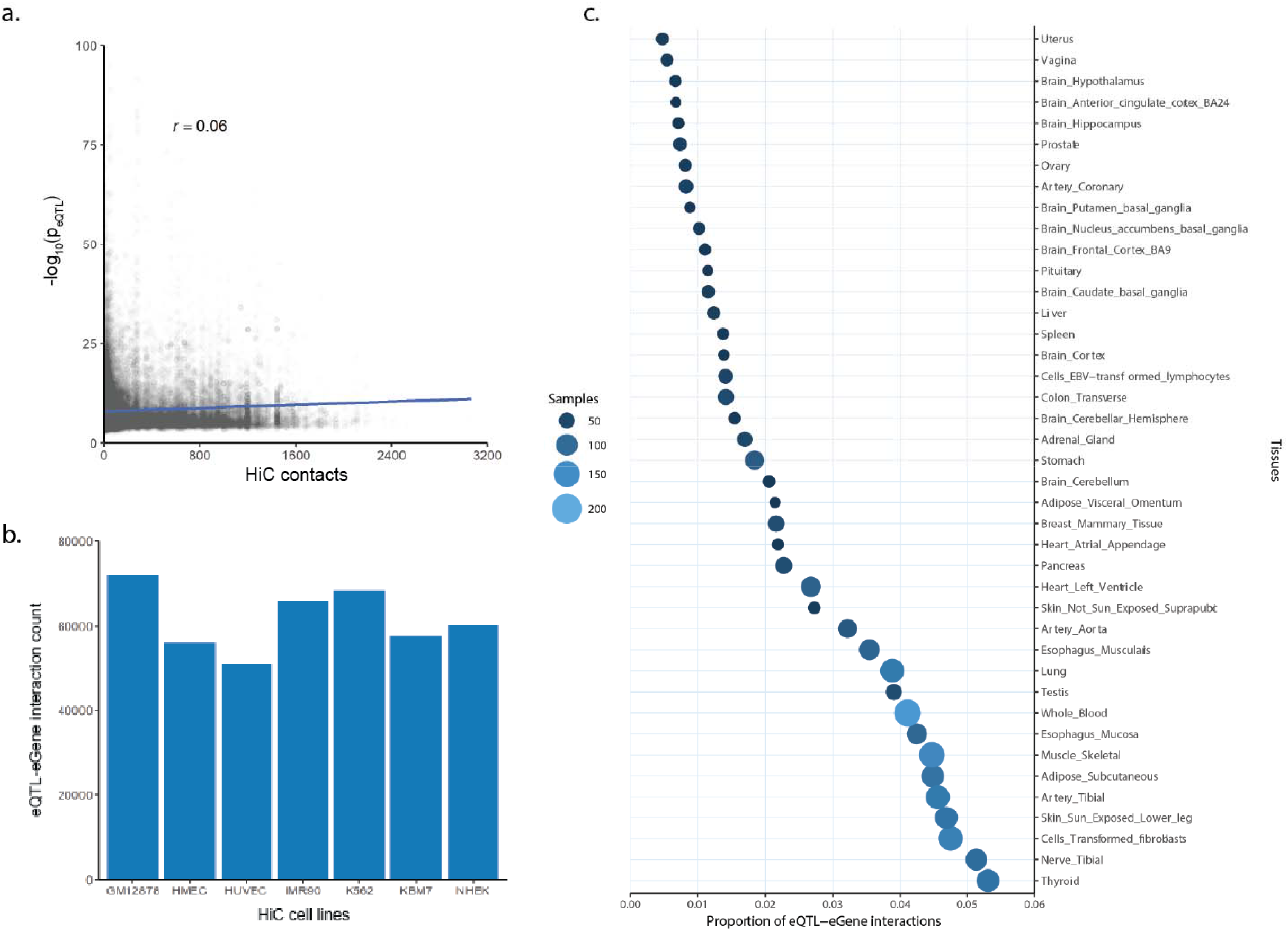
Tissue specificity of spatial eQTL-eGene interactions. (a) A plot of all significant eQTL *p* values on eGene against the number of HiC contacts for the interactions. (b) Distribution of eQTL-eGene interactions among the HiC cell lines. (c) The proportion of eQTL-eGene interactions in tissues positively correlates (*r* = 0,87) with the number of RNASeq and genotyped samples in GTEx.

## Tables List

Supplementary Tables are available at doi.orq/<u>10.17608/k6.auckland.6291728</u>

**S1 Table. eGene interactions**.

**S2 Table. Shared eGene matrix**.

**S3 Table. Clusters of phenotypes**.

**S4. eQTLs in the fat metabolism cluster**.

